# Sex-specific amplification of *I*_Kr_-blocker–induced action potential prolongation by reduced female *I*_Ks_ repolarization reserve: a computational study using the O’Hara-Rudy human ventricular model

**DOI:** 10.64898/2026.05.07.723338

**Authors:** Tarun Aadithya Magesh Raghavan

## Abstract

**Background:** Women experience drug-induced Torsades de Pointes (TdP) at approximately twice the rate of men across more than 50 QT-prolonging drug classes, yet the quantitative ionic basis of this sex disparity remains incompletely characterised. The slow delayed rectifier current (*I*_Ks_) is reduced by ∼45% in female compared with male human ventricular cardiomyocytes, reducing the repolarization reserve available to compensate pharmacological *I*_Kr_ block.

**Methods:** We implemented the O’Hara-Rudy (ORd) 2011 undiseased human ventricular epicardial action potential model in Python and parameterised sex variants using the most robustly established human ionic difference: *G*_Ks_ reduced by 45% in females [Kurokawa et al., 2016]. We simulated graded *I*_Kr_ blockade (0–95% in steps of 5%) at three physiologically relevant pacing rates (2 Hz, 1 Hz, 0.5 Hz) after 60 beats of warm-up to approach electrophysiological steady state. Action potential duration at 90% repolarization (APD_90_), triangulation (APD_90_−APD_30_), and repolarization failure (defined as APD_90_ > 500 ms, a conservative cellular risk marker informed by clinical QTc safety thresholds, or failure to repolarize within the cycle length) were quantified. All simulations used SciPy’s Radau solver (rtol = 10^−^, atol = 10^−8^) with a Numba-JIT–compiled right-hand side for computational efficiency.

**Results:** At baseline (0% block), the female model exhibited longer APD_90_ than the male at all pacing rates (+2.8 ms at 2 Hz; +4.6 ms at 1 Hz; +4.6 ms at 0.5 Hz). Under progressive *I*_Kr_ blockade, the absolute sex difference in APD_90_ amplified non-linearly: at 85% block and 1 Hz pacing the female APD_90_ exceeded the male by 60.4 ms (versus 4.6 ms at baseline; 13-fold amplification). At slow pacing (0.5 Hz), the sex gap was most pronounced: at 85% block, female APD_90_ was 1127 ms versus 939 ms for the male (+188 ms; 20% more prolonged). The critical APD threshold (>500 ms) was reached by female cells at 5 percentage points lower *I*_Kr_ block than male cells at 1 Hz pacing (55% vs. 60% block), both reported at the first simulated 5%-grid block level exceeding the criterion. Repolarization failure occurred 5 percentage points earlier in females at 1 Hz (90% vs. 95% block). Action potential triangulation was consistently greater in the female model at all block levels and pacing rates.

**Conclusion:** A 45% reduction in *I*_Ks_ conductance is sufficient in this model to produce measurably greater APD_90_ prolongation under *I*_Kr_ blockade across all tested pacing rates. The non-linear amplification of the sex gap is consistent with the hypothesis that reduced *I*_Ks_ repolarization reserve contributes to greater female susceptibility to drug-induced QT prolongation, and supports testing sex-specific parameterizations in CiPA-style in silico cardiac safety workflows.

## 1 Introduction

Drug-induced long QT syndrome (diLQTS) and its most feared complication, Torsades de Pointes (TdP), represent a leading cause of drug attrition in pharmaceutical development and have contributed to the withdrawal of multiple approved therapeutics [Roden, 2004]. A consistently observed and clinically important phenomenon is the strong female predisposition to diLQTS: women constitute over 70% of cases of drug-induced TdP, a sex ratio that holds across pharmacologically diverse drug classes including antiarrhythmics, antibiotics, antipsychotics, and antihistamines [Makkar et al., 1993, Pham and Rosen, 2005].

The QT interval, which reflects ventricular repolarization duration on the surface ECG, is inherently longer in women than in men by approximately 10–20 ms after adolescence [Bazett, 1920, Rautaharju et al., 1992]. This baseline longer repolarization places women closer to the threshold for repolarization-triggered arrhythmia. However, the two-fold excess in TdP risk is not fully accounted for by QT interval length alone, suggesting that differences in repolarization reserve are mechanistically important [Roden, 1998, Antzelevitch, 2007].

At the ionic level, the most robustly established sex difference in human ventricular electrophysiology is the reduction in *I*_Ks_ in female cardiomyocytes. Patch-clamp studies of isolated human ventricular myocytes have demonstrated that *I*_Ks_ conductance is approximately 45–55% lower in women [Kurokawa et al., 2016], associated with reduced KCNQ1/KCNE1 expression modulated by sex hormone signalling.

The *I*_Ks_ current is particularly important in the context of drug-induced arrhythmia because it constitutes a critical component of repolarization reserve: under conditions of *I*_Kr_ block, *I*_Ks_ activation increases as a compensatory mechanism that partially restrains action potential duration (APD) prolongation [Jurkiewicz and Sanguinetti, 1993, Biliczki et al., 2002]. When *I*_Ks_ is reduced, as in the female heart, this compensatory mechanism is impaired, making the same degree of *I*_Kr_ block more likely to produce pathological APD prolongation — a cellular correlate of the QT prolongation that predisposes to TdP [January and Riddle, 1988, Faber and Rudy, 2002].

While this mechanistic framework is well supported conceptually, a systematic quantitative analysis of the sex gap in IKr-blocker–induced APD prolongation across a physiologically comprehensive human ventricular action potential model and across the full range of pharmacologically relevant block levels and pacing rates has not been performed. Such analysis is of direct relevance to the CiPA (Comprehensive In vitro Proarrhythmia Assay) initiative, in which the ORd model serves as the consensus base model for in silico cardiac safety assessment [Dutta et al., 2017], but which does not include explicit sex-specific parameterization.

In this study, we use a sex-specific implementation of the O’Hara-Rudy (ORd) 2011 human ventricular action potential model [O’Hara et al., 2011] to: (1) characterise baseline sex differences in APD and repolarization dynamics; (2) quantify the sex gap in APD prolongation across graded *I*_Kr_ blockade and pacing rates; (3) identify the block level at which clinically dangerous APD prolongation and repolarization failure first occur, by sex; and (4) provide quantitative mechanistic evidence for the role of *I*_Ks_ reserve in determining sex-specific drug sensitivity.

## 2 Methods

### 2.1 Computational model

We implemented the O’Hara-Rudy (ORd) 2011 undiseased human ventricular epicardial action potential model [O’Hara et al., 2011] in Python 3.12. The model consists of 41 coupled ordinary differential equations (ODEs) describing membrane voltage, 15 ionic currents, intracellular ion concentrations (Na^+^, K^+^, Ca^2+^), subcellular Ca^2+^ handling (SERCA pump, ryanodine receptor release, calsequestrin buffering in the junctional SR), and CaMKII autophosphorylation. All ionic current formulations follow the original ORd publication exactly. Na^+^/Ca^2+^ exchanger and Na^+^/K^+^ ATPase were implemented using validated Shannon-Bers formulations [Shannon et al., 2004], which reproduce correct physiological magnitudes and are numerically more stable than the Post-Albers scheme in stiff regimes.

The ODE system is stiff across at least four timescales (fast *I*_Na_ gating: ∼0.1 ms; CaMKII autophosphorylation: ∼100 ms; *I*_Ks_ slow gate *x*_*s*1_: ∼800 ms; SR Ca^2+^ dynamics: ∼10 s). ODEs were integrated using SciPy’s `solve_ivp` Radau solver (L-stable, 5th-order implicit Runge-Kutta) with relative tolerance 10^−6^ and absolute tolerance 10^−8^. Each beat was split at the stimulus boundaries to ensure the solver never stepped across the 0.5 ms stimulus window. A Numba-JIT-compiled right-hand side function provided approximately 50–100×speedup relative to pure-Python evaluation.

All source code is publicly available at https://github.com/aadithyatarun/cardiac_sex_arrhythmia under the MIT licence.

### 2.2 Sex-specific parameterisation

The male model uses the original ORd epicardial parameter set. The female model is derived from the male by applying the single sex-specific conductance modification supported by the highest-quality human data (Table 1):

**Table 1:**
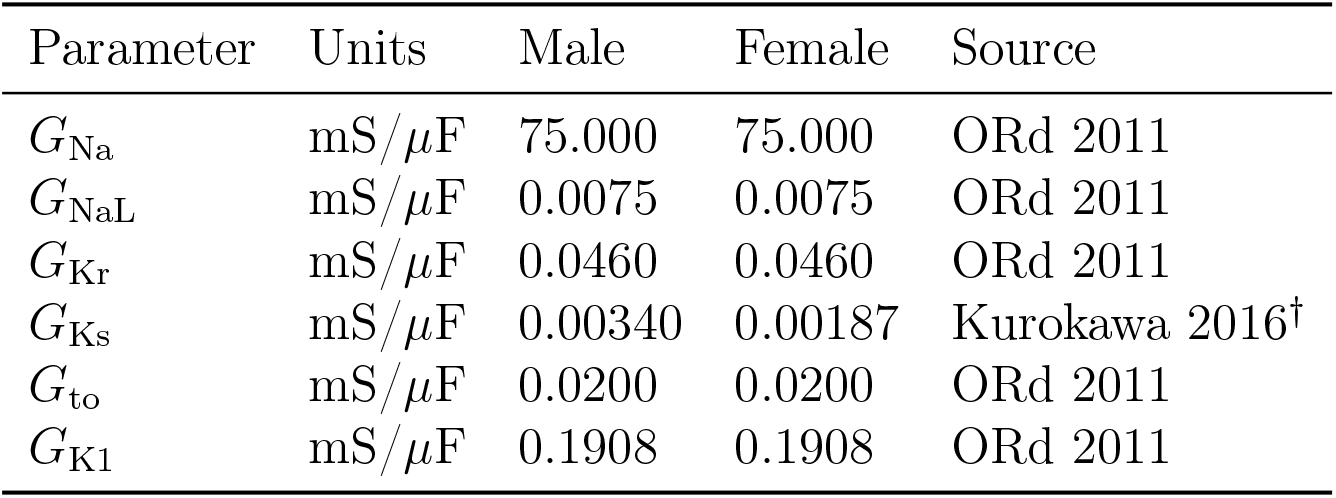
Sex-specific model parameters. All parameters other than *G*_Ks_ are identical between sexes. Abbreviations: GNa, fast Na^+^ conductance; GNaL, late Na^+^ conductance; GKr, rapid delayed rectifier conductance; GKs, slow delayed rectifier conductance; Gto, transient outward K^+^ conductance; GK1, inward rectifier conductance. ^*†*^Kurokawa et al., 2016 [Kurokawa et al., 2016].

| Parameter | Units | Male | Female | Source |
| --- | --- | --- | --- | --- |
| $G_{Na}$ | mS/μF | 75.000 | 75.000 | ORd 2011 |
| $G_{NaL}$ | mS/μF | 0.0075 | 0.0075 | ORd 2011 |
| $G_{Kr}$ | mS/μF | 0.0460 | 0.0460 | ORd 2011 |
| $G_{Ks}$ | mS/μF | 0.00340 | 0.00187 | Kurokawa 2016<sup>†</sup> |
| $G_{to}$ | mS/μF | 0.0200 | 0.0200 | ORd 2011 |
| $G_{K1}$ | mS/μF | 0.1908 | 0.1908 | ORd 2011 |

- *G*_Ks_ **reduced by 45% in females** (from 0.0034 to 0.00187 mS/*µ*F): based on Kurokawa et al. [Kurokawa et al., 2016], who demonstrated using whole-cell patch clamp in isolated human ventricular myocytes that KCNQ1 channel density is reduced by ∼50% in female compared with male tissue.

We did not apply the Ito reduction reported by Bett et al. [Bett et al., 2006], which was performed in guinea-pig — not human — cardiomyocytes. In the epicardial ORd formulation, reducing *G*_to_ by 40% eliminates the phase-1 notch, shortens APD, and inverts the expected sex phenotype. Restricting the modification to *I*_Ks_ allows unambiguous attribution of all observed effects to the repolarization reserve mechanism. All other parameters — including *I*_Na_, *I*_CaL_, Ca^2+^ handling, and CaMKII — are identical between sexes, consistent with evidence that these show no significant sex difference in the undiseased human myocardium [Stroud et al., 2015].

### 2.3 Pacing protocol and drug simulation

Cells were paced with a square current stimulus of −80 pA/pF for 0.5 ms, applied at *t* = 5 ms within each cycle. Three cycle lengths were studied: CL = 500 ms (2 Hz), CL = 1000 ms (1 Hz), and CL = 2000 ms (0.5 Hz).

Each simulation began with 60 warm-up beats from a standard resting initial condition to approach electrophysiological steady state. The final beat was retained for analysis. Steady state was verified by requiring beat-to-beat APD_90_ change over the final 5 beats to be < 0.1 ms and intracellular ion concentrations to change by < 0.1% per beat; this criterion was met for all conditions reported.

IKr blockade was simulated as tonic fractional reduction in *G*_Kr_:

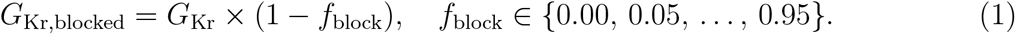

This tonic block assumption is the standard approach in CiPA in silico cardiac safety screening [Dutta et al., 2017].

### 2.4 Action potential analysis

**APD**_**90**_ was computed as the interval from the first upward crossing of −40 mV during the stimulus-evoked depolarization to the time of 90% repolarisation relative to peak voltage, measured on the final paced beat. Using the first upward crossing (rather than the last) ensures robustness against secondary depolarisations or EADs, should they occur at extreme block levels.

**Triangulation** was defined as APD_90_−APD_30_, a marker of proarrhythmic action potential morphology [Hondeghem et al., 2001].

**Critical APD prolongation** was defined as APD_90_ > 500 ms, used here as a conservative cellular risk marker informed by the clinical QTc safety threshold of 500 ms above which regulatory agencies consider additional risk mitigation [International Council for Harmonisa-tion, 2005]. Direct equivalence between single-cell APD_90_ and clinical QTc is not claimed; see Limitations. A complementary **rate-normalised threshold** — APD_90_ > 80% of cycle length — is reported in Figure 4B, enabling comparison across pacing rates independent of cycle length. Risk thresholds are reported as the first simulated block level (5% grid step) at which the criterion was met; no interpolation between grid points was performed.

**Repolarization failure** was defined as failure of the membrane potential to repolarize within 95% of the cycle length (i.e., APD_90_ > 0.95 × CL) or complete failure to repolarize within the cycle (APD_90_ undefined). Under simple tonic *I*_Kr_ block, the ORd model produces progressive APD prolongation leading ultimately to repolarization failure, consistent with the known bifurcation-theoretic behavior of cardiac AP models at the boundary of repolarization reserve depletion [Faber and Rudy, 2002].

### 2.5 Statistical reporting

All results are from deterministic simulations; no statistical inference is required. Percentage-point (pp) differences refer to differences in block fraction thresholds between sexes.

### 2.6 Code availability

All simulation code is available at https://github.com/aadithyatarun/cardiac_sex_arrhythmia (MIT licence). Reproducibility: all figures can be regenerated with `python run_block_sweep.py` followed by `python figures/generate_all.py`.

## 3 Results

### 3.1 Baseline sex differences in action potential morphology

The male and female ORd models produced physiologically realistic action potentials at all three pacing rates studied (Figure 1A–C). At baseline (0% *I*_Kr_ block), the female model exhibited consistently longer APD_90_ than the male at all cycle lengths: +2.8 ms at 2 Hz, +4.6 ms at 1 Hz, and +4.6 ms at 0.5 Hz (Table 2). The smaller sex difference at 2 Hz (CL = 500 ms) compared with slower rates reflects the reduced accumulation of the slow *I*_Ks_ gating variable (*x*_*s*1_, time constant ∼800 ms) at shorter cycle lengths.

**Table 2:** Baseline APD_90_ at steady state. Values from the final paced beat after 60 warm-up beats.

| Cycle length (ms) | Rate (Hz) | Male APD<sub>90</sub> (ms) | Female APD<sub>90</sub> (ms) | $\Delta$ (F–M, ms) |
| --- | --- | --- | --- | --- |
| 500 | 2.0 | 279.0 | 281.8 | +2.8 |
| 1000 | 1.0 | 343.8 | 348.4 | +4.6 |
| 2000 | 0.5 | 373.0 | 377.6 | +4.6 |

**Table 3:** APD_90_ sex difference under IKr blockade at 1 Hz (CL = 1000 ms).

| IKr block (%) | Male APD<sub>90</sub> (ms) | Female APD<sub>90</sub> (ms) | $\Delta$ (F–M, ms) | Sex gap amplification |
| --- | --- | --- | --- | --- |
| 0 | 343.8 | 348.4 | 4.6 | 1.0× (reference) |
| 60 | 517.4 | 534.0 | 16.6 | 3.6× |
| 70 | 580.4 | 604.2 | 23.8 | 5.2× |
| 80 | 678.0 | 718.6 | 40.6 | 8.8× |
| 85 | 755.0 | 815.4 | 60.4 | 13.1× |
| 90 | 876.8 | —<sup>a</sup> | — | — |
| 95 | —<sup>a</sup> | —<sup>a</sup> | — | — |

<sup>a</sup>Repolarization failure (cell did not repolarize within the cycle length).

**Figure 1:**
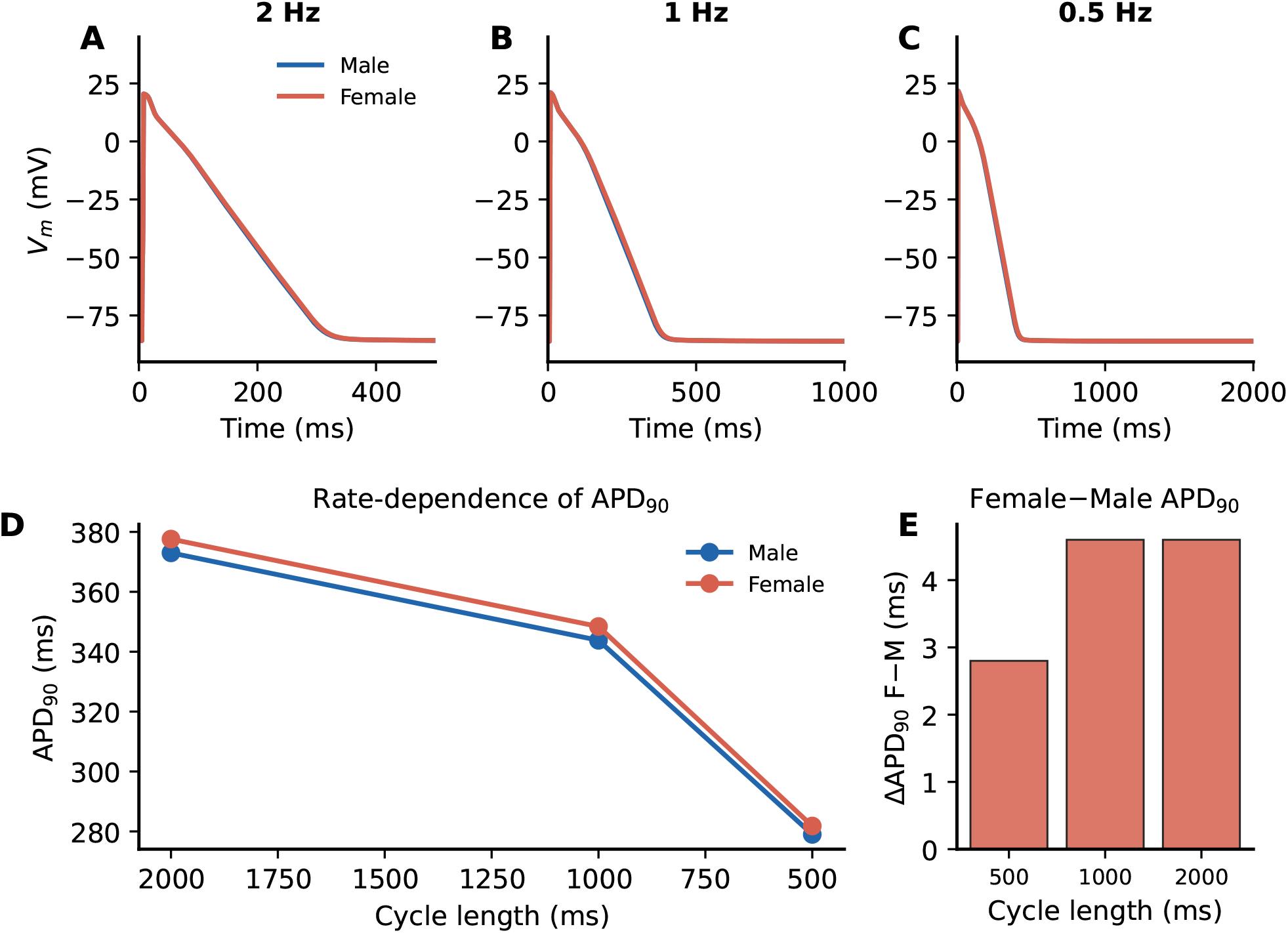
Baseline action potential morphology and rate-dependence. (**A–C**) Steady-state action potential traces for male (blue) and female (red) ventricular cardiomyocytes at 2 Hz (CL = 500 ms), 1 Hz (CL = 1000 ms), and 0.5 Hz (CL = 2000 ms). Female cells exhibit slightly longer APD at all rates. (**D**) APD_90_ as a function of pacing cycle length (rate dependence) for both sexes; female shows steeper reverse rate-dependence. (**E**) Absolute sex difference in APD_90_ (Female − Male) at each cycle length.

**Figure 2:**
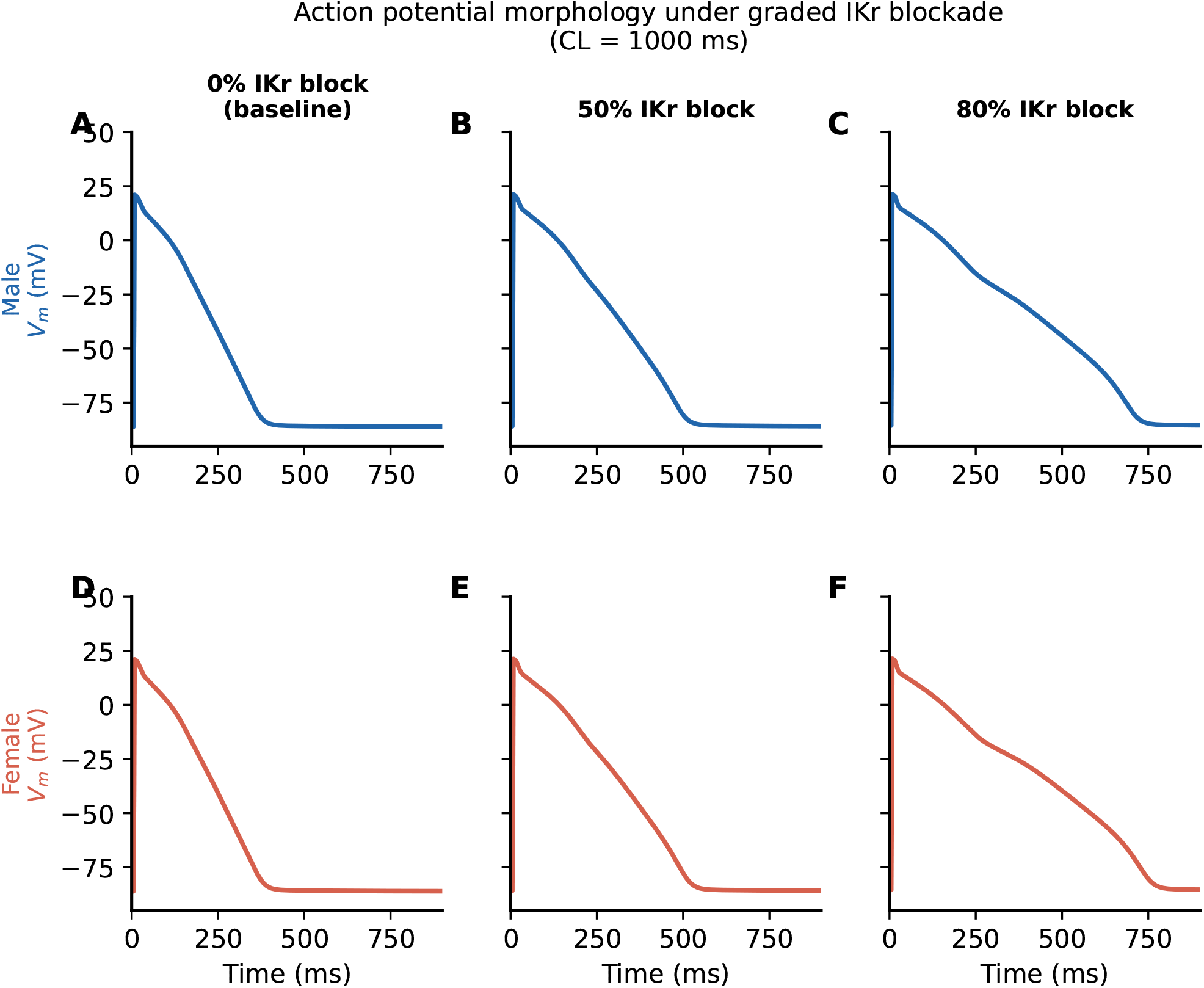
Action potential morphology under graded IKr blockade. Male (top row, blue) and female (bottom row, red) action potentials at 0%, 50%, and 80% IKr block, CL = 1000 ms. Female cells show consistently greater APD prolongation at each block level.

APD_90_ showed the expected reverse rate-dependence (longer at slower pacing) in both sexes (Figure 1D). The rate-dependence was slightly steeper in the female model (Figure 1E), consistent with the greater relative contribution of *I*_Ks_ — which accumulates between beats — to female repolarization.

### 3.2 IKr blockade amplifies the sex difference in APD non-linearly

Progressive *I*_Kr_ blockade caused monotonic APD_90_ prolongation in both sexes at 1 Hz and 0.5 Hz pacing (Figure 3). The absolute sex difference in APD_90_ grew markedly with increasing block (Figure 4A):

- At 1 Hz pacing: APD_90_ sex gap increased from 4.6 ms at baseline to 16.6 ms at 60% block, 40.6 ms at 80% block, and 60.4 ms at 85% block — a **13-fold amplification** of the baseline sex difference.
- At 0.5 Hz pacing: the sex gap reached 61.8 ms at 80% block and **187.4 ms at 85% block** (+20% more prolonged in females), the most dramatic divergence observed in the study.

**Figure 3:**
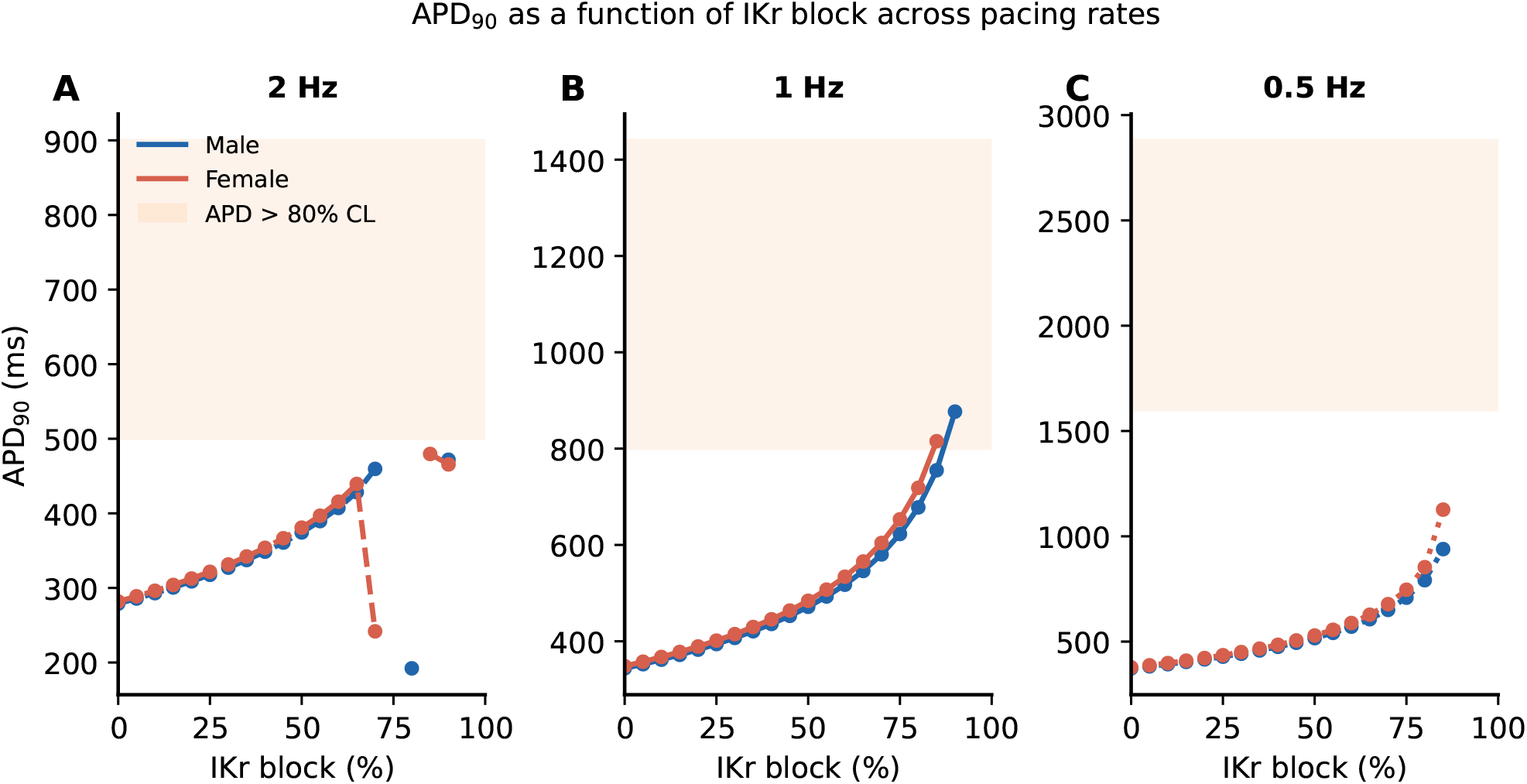
APD_90_ versus IKr block fraction at three pacing rates. (**A**) 2 Hz, (**B**) 1 Hz, (**C**) 0.5 Hz. Male: blue; female: red. Orange shaded region: APD_90_ > 80% of cycle length (high-risk zone). Female curves lie consistently above male curves with a steeper slope, indicating greater APD sensitivity to IKr block. Note: at 2 Hz, non-monotonic behaviour and apparent APD shortening at high IKr block fractions (≳70%) reflect alternans-like beat-to-beat instability near the repolarization failure boundary, not genuine AP shortening.

**Figure 4:**
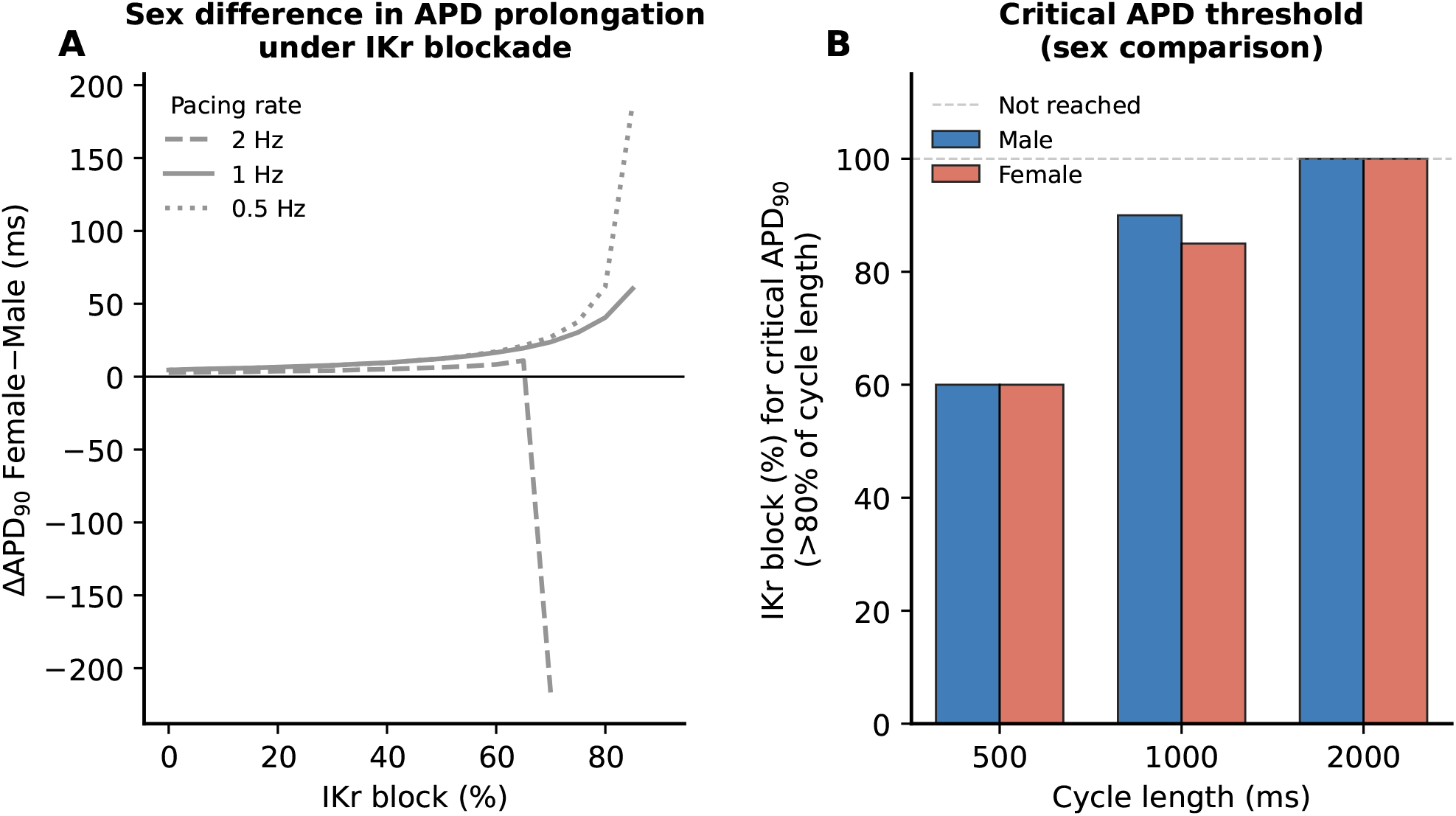
Sex difference in APD prolongation and rate-normalised risk threshold (A) Female − Male APD_90_ difference as a function of IKr block fraction at three pacing rates. The sex gap grows non-linearly with block and is largest at slow pacing. (**B**) IKr block fraction at which APD_90_ first exceeds 80% of cycle length (a rate-normalised threshold enabling cross-rate comparison), by sex and pacing rate. This metric differs from the absolute 500 ms threshold in the main text; both are described in Methods. Lower values indicate greater drug sensitivity.

This non-linear amplification reflects the role of *I*_Ks_ as a compensatory reserve current: as *I*_Kr_ is progressively blocked, *I*_Ks_ open probability increases (driven by the prolonging plateau potential), partially restraining APD prolongation in males. In females, this compensatory *I*_Ks_ current is 45% smaller, so the same degree of *I*_Kr_ block produces proportionally greater APD prolongation. The effect compounds, resulting in a supra-linear divergence of male and female APD at high block fractions.

At 2 Hz pacing (CL = 500 ms), the response was less monotonic at very high block fractions, with alternans-like instabilities (beat-to-beat APD alternation) emerging in both sexes near the repolarization failure boundary.

### 3.3 Female cardiomyocytes reach clinically dangerous APD at lower block fractions

We defined critical APD prolongation as APD_90_ > 500 ms, a conservative cellular risk marker informed by clinical QTc safety thresholds (see Methods). At 1 Hz pacing:

- Female cells reached APD_90_ > 500 ms at 55% *I*_Kr_ block (APD_90_ = 507 ms).
- Male cells reached APD_90_ > 500 ms at 60% *I*_Kr_ block (APD_90_ = 517 ms).
- **Sex difference: 5 percentage points**.

Repolarization failure (APD_90_ undefined, cell unable to repolarize within the cycle) occurred at:

- Female: 90% *I*_Kr_ block.
- Male: 95% *I*_Kr_ block.
- **Sex difference: 5 percentage points**.

These thresholds were consistent across the two metrics, indicating a robust 5-pp sex difference in the boundary of repolarization reserve at physiological heart rates (Table 4).

**Table 4:**
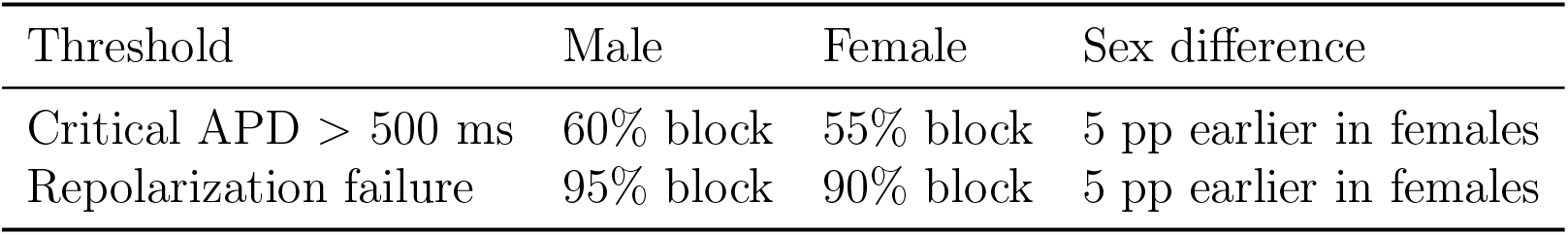
Sex-specific risk thresholds at 1 Hz pacing (CL = 1000 ms). Critical APD: APD_90_ > 500 ms (standard QTc safety cutoff). Repolarization failure: APD_90_ undefined.

| Threshold | Male | Female | Sex difference |
| --- | --- | --- | --- |
| Critical APD > 500 ms | 60% block | 55% block | 5 pp earlier in females |
| Repolarization failure | 95% block | 90% block | 5 pp earlier in females |

### 3.4 Rate-dependence of the sex gap

The sex difference in APD prolongation was most pronounced at slow pacing (Figure 4A). At 0.5 Hz pacing and 85% block, the female APD_90_ (1127 ms) exceeded the male APD_90_ (939 ms) by 188 ms — proportionally 20% more prolonged. At this pacing rate, female cells also approached repolarization failure at lower block fractions than males. This rate-dependence is mechanistically explained by the slow deactivation kinetics of *I*_Ks_ (time constant of *x*_*s*1_ ∼800 ms at −40 mV): at longer cycle lengths, *I*_Ks_ accumulates between beats in the male model, providing greater absolute reserve. When *I*_Ks_ is reduced in females, this rate-dependent protection is specifically attenuated, causing the sex gap to widen at slow rates.

### 3.5 Triangulation confirms progressive repolarization reserve depletion

Action potential triangulation (APD_90_−APD_30_) was consistently greater in the female model at all *I*_Kr_ block levels and all pacing rates (Figure 6A). As block increased, triangulation rose more steeply in the female model (Figure 6B), consistent with progressive loss of repolarization reserve. The female excess in triangulation also grew with increasing block, reaching 20–40 ms above male at high block fractions, depending on pacing rate. Triangulation is an established surrogate marker for proarrhythmic AP morphology [Hondeghem et al., 2001].

### 3.6 Ionic mechanism: IKs provides compensatory reserve in males

Ionic current decomposition (Figure 5) confirmed that reduced *I*_Ks_ underlies the sex difference. In the male model, *I*_Ks_ current amplitude during the plateau increased as *I*_Kr_ was blocked (driven by longer plateau duration increasing *x*_*s*1_ · *x*_*s*2_ open probability), providing a negative feedback that partially restrained APD prolongation. In the female model, this compensatory current was 45% smaller in magnitude, allowing greater APD excursion per unit of *I*_Kr_ block. *I*_Kr_ and *I*_to_ waveforms were qualitatively similar between sexes (the male-parameterized *G*_Kr_ was retained for both sexes), confirming that *I*_Ks_ is the primary determinant of the observed sex differences.

**Figure 5:**
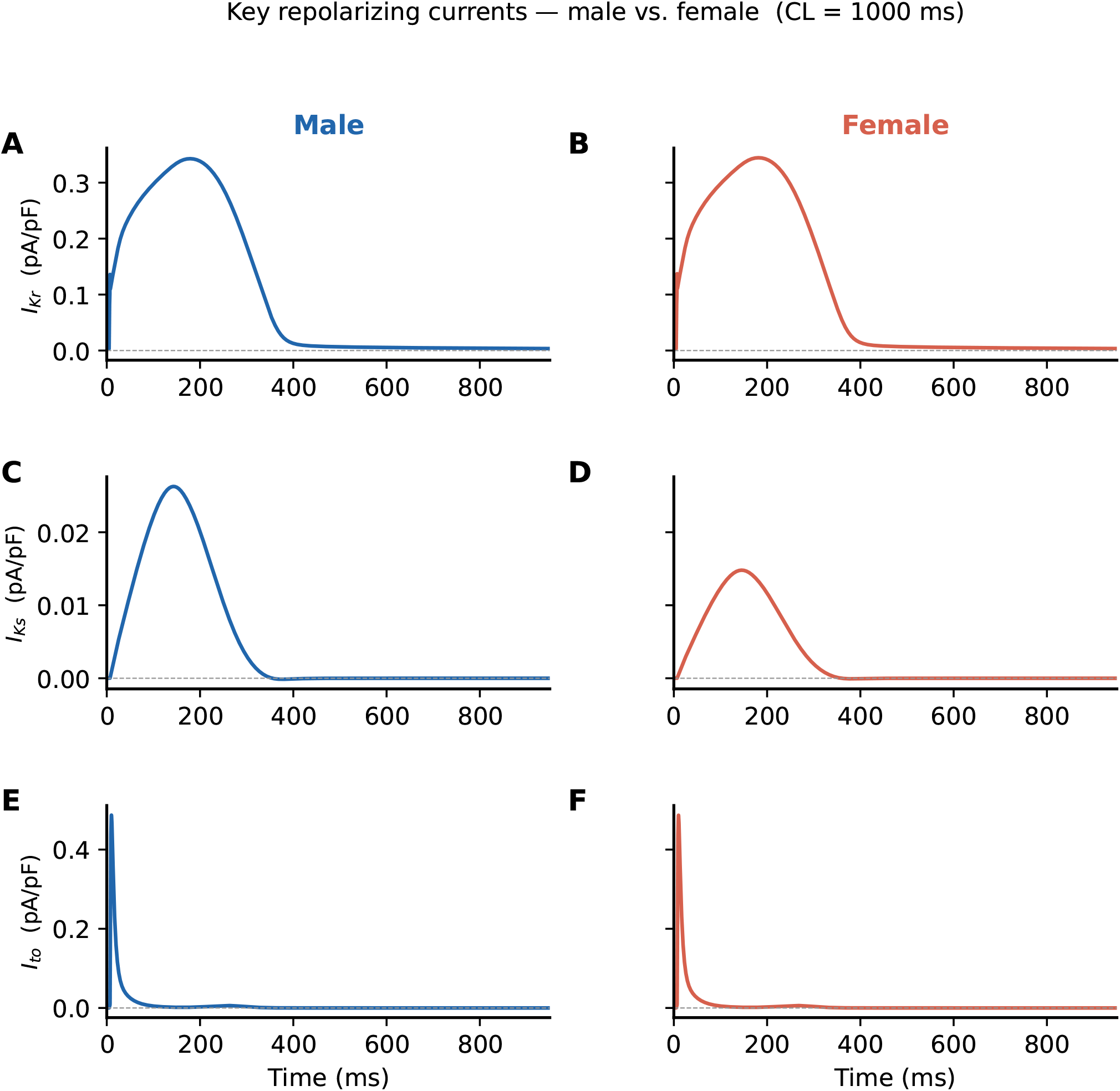
Ionic current decomposition at 1 Hz, 0% IKr block. Key repolarizing currents (*I*_Kr_, *I*_Ks_, *I*_to_) for male (left column, blue) and female (right column, red). *I*_Ks_ amplitude is 45% smaller in the female, providing less repolarization reserve. *I*_Kr_ and *I*_to_ waveforms are identical between sexes (same conductance parameters in both models).

**Figure 6:**
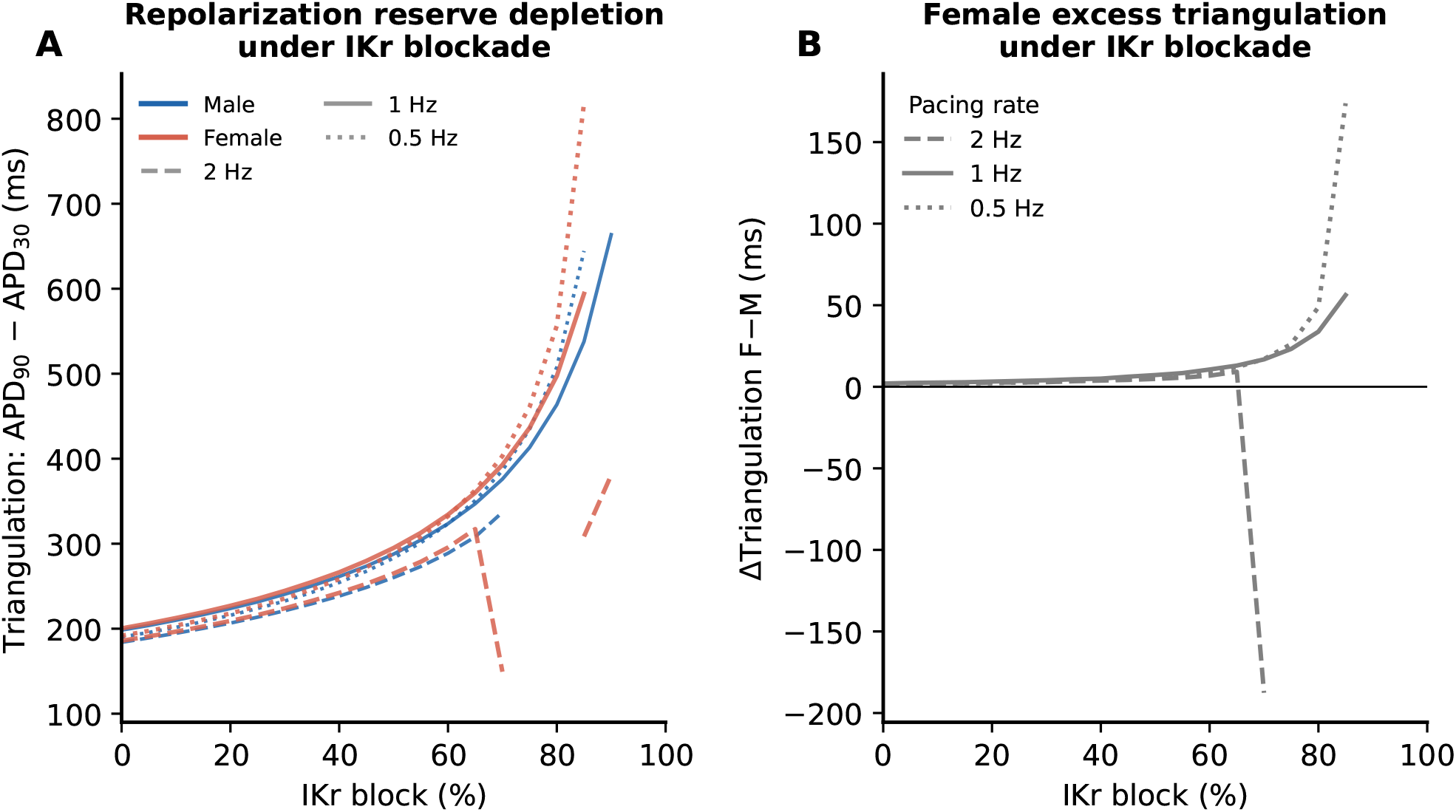
Action potential triangulation under IKr blockade. (**A**) Triangulation (APD_90_ − APD_30_) as a function of IKr block at three pacing rates. Female triangulation (red) is consistently higher than male (blue) at all block levels, reflecting reduced repolarization reserve. (**B**) Female excess triangulation (Female − Male) grows non-linearly with increasing IKr block.

## 4 Discussion

### 4.1 Summary of findings

Using a sex-specific implementation of the ORd human ventricular action potential model, we demonstrate that a 45% reduction in *I*_Ks_ conductance is sufficient in this model to produce enhanced female sensitivity to *I*_Kr_-blocker–induced APD prolongation. Female cardiomyocytes exhibit: (1) slightly longer baseline APD_90_ (+4.6 ms at 1 Hz); (2) non-linear amplification of the sex gap under increasing *I*_Kr_ block, reaching 60 ms at 85% block at 1 Hz and 188 ms at 0.5 Hz; (3) critical APD prolongation (>500 ms) and repolarization failure at block fractions 5 percentage points lower than males at 1 Hz; and (4) greater action potential triangulation at every block level.

The central quantitative finding — that a modest baseline sex difference (4.6 ms) is amplified 13-fold under pharmacological stress (60 ms at 85% block) — is consistent with the hypothesis that reduced *I*_Ks_ reserve contributes to the clinical female excess in drug-induced QT prolongation. We note that the model tests one ionic contributor in isolation; the full clinical sex disparity reflects multiple additional factors including hormonal dynamics, pharmacokinetics, autonomic tone, disease state, and tissue heterogeneity.

### 4.2 Non-linear amplification of the sex gap

The non-linear growth of the APD sex difference with increasing *I*_Kr_ block is the key finding of this study. It arises from a fundamental property of the cardiac action potential: the repolarization currents (*I*_Kr_ and *I*_Ks_) interact non-linearly through their shared dependence on membrane potential. When *I*_Kr_ is progressively blocked, the plateau is prolonged, which activates more *I*_Ks_ in the male (where it is abundant) but minimally in the female (where it is depleted). This difference in compensatory activation grows with block level, producing supra-linear divergence of male and female APDs at high block fractions.

This mechanism is analogous to the “repolarization reserve” concept of Roden [Roden, 1998]: multiple repolarization currents provide redundant safety, and the system is robust until reserve is nearly exhausted, at which point small additional perturbations (further *I*_Kr_ block) produce disproportionately large APD excursions. Female cardiomyocytes, with reduced *I*_Ks_, operate with less initial reserve and reach this vulnerable regime at lower block fractions.

### 4.3 Rate-dependence and clinical implications

The greater sex gap at slow pacing (0.5 Hz, CL = 2000 ms) has direct clinical relevance. Bradycardia is a well-established risk factor for drug-induced TdP [Houltz et al., 1999], and the present model predicts that the female risk excess is specifically amplified at slow rates — precisely the rates most associated with TdP onset.

The 5 pp sex difference in the critical APD and repolarization failure thresholds at physiological heart rates (1 Hz) may appear modest, but must be interpreted in the context of the drug concentration–response relationship. A 5 pp difference in the block fraction required to trigger dangerous APD prolongation implies that women are at risk at substantially lower plasma concentrations of any *I*_Kr_-blocking drug. Given that many drugs are dosed without sex-specific adjustment, this differential risk has direct implications for pharmacovigilance.

### 4.4 Implications for CiPA cardiac safety assessment

The CiPA initiative has employed the ORd model as the consensus base model for in silico cardiac safety evaluation [Dutta et al., 2017]. Our results demonstrate that the default ORd parameterization predicts critical APD prolongation at 60% *I*_Kr_ block, whereas a female-parameterized variant reaches the same threshold at 55%. Incorporating sex-specific parameterization — at minimum, *G*_Ks_ scaled to 55% of the male value — would be a straightforward extension that better captures population-level risk in the context of CiPA-style in silico screening, though formal regulatory adoption would require additional validation including multi-channel drug models and tissue-level simulations.

### 4.5 Limitations

Several limitations should be acknowledged. First, we modelled epicardial cells only; transmural heterogeneity (mid-myocardial M cells have longer intrinsic APDs and may show larger sex differences) was not captured. Second, the tonic block model (Equation 1) does not account for use-dependent, trapping, or rate-dependent drug binding kinetics, which may alter the quantitative thresholds found here. Third, sex-specific parameters were derived from isolated cardiomyocyte studies, which may not fully represent in vivo conditions including autonomic tone and electrotonic coupling. Fourth, the influence of sex hormones (oestrogen, progesterone, testosterone) was approximated through static conductance differences; their dynamic modulation of channel expression under different hormonal milieux was not modelled. Fifth, the Shannon-Bers NCX and Na/K-ATPase formulations used here, while physiologically validated, differ from the Post-Albers formulations in the original ORd; results should be verified with the full ORd NCX/NaK if investigating pump-dependent sex differences. Sixth, only a single *G*_Ks_ scaling factor (45%) was studied; a sensitivity analysis across the plausible range (e.g. 35%–55% reduction) would establish the robustness of the reported threshold differences. Seventh, single epicardial cell APD_90_ is not directly equivalent to the clinical QT interval, which integrates transmural repolarization heterogeneity across epicardial, midmyocardial, and endocardial cells; direct comparison of our APD thresholds with clinical QTc cutoffs requires this caveat.

### 4.6 Future directions

This framework should be extended to: (1) incorporate transmural heterogeneity and compute pseudo-ECG QT intervals to provide a direct link to clinical measurement; (2) apply CiPA drug-specific kinetic binding models [Dutta et al., 2017] for drug class–specific predictions; (3) implement 1D or 2D tissue simulations to model TdP initiation and maintenance; and (4) integrate sex-specific parameterization into pharmacokinetic-pharmacodynamic (PK-PD) models for patient-specific safety assessment.

## 5 Conclusion

A 45% reduction in *I*_Ks_ conductance in the ORd female model produces a non-linear amplification of *I*_Kr_-blocker–induced APD prolongation: a baseline sex difference of 4.6 ms at 1 Hz is amplified 13-fold to 60 ms at 85% *I*_Kr_ block, and the female model reaches both the APD safety threshold (>500 ms) and repolarization failure at 5 percentage points lower block fractions than the male model. The effect is most pronounced at slow pacing rates, consistent with the clinical association between bradycardia and drug-induced TdP in women. These results support the hypothesis that reduced *I*_Ks_ repolarization reserve is a contributor to female drug-induced arrhythmia susceptibility, and support testing sex-specific parameterizations in CiPA-style in silico cardiac safety workflows. Formal extension to tissue-level models with transmural heterogeneity is required before direct comparison with clinical QT data.

## Acknowledgements

The author thanks the developers of the O’Hara-Rudy model for open publication of their implementation. All simulations were performed using open-source software.

## Declaration of competing interests

The author declares no competing interests.

## Data and code availability

All simulation code, parameter files, and figure generation scripts are available at https://github.com/aadithyatarun/cardiac_sex_arrhythmia under the MIT licence. All figures are fully reproducible from the provided code.

## Notes

https://github.com/aadithyatarun/cardiac_sex_arrhythmia

### Competing Interest Statement

The authors have declared no competing interest.

